# Geological processes mediate a subsurface microbial loop in the deep biosphere

**DOI:** 10.1101/2021.10.26.465990

**Authors:** Daniel A. Gittins, Pierre-Arnaud Desiage, Natasha Morrison, Jayne E. Rattray, Srijak Bhatnagar, Anirban Chakraborty, Jackie Zorz, Carmen Li, Oliver Horanszky, Margaret A. Cramm, Jamie Webb, Adam MacDonald, Martin Fowler, D. Calvin Campbell, Casey R. J. Hubert

## Abstract

The deep biosphere is the largest microbial habitat on Earth and features abundant bacterial endospores^1,2^. Whereas dormancy and survival at theoretical energy minima are hallmarks of subsurface microbial populations^3^, the roles of fundamental ecological processes like dispersal and selection in these environments are poorly understood^4^. Here we combine geophysics, geochemistry, microbiology and genomics to investigate biogeography in the subsurface, focusing on bacterial endospores in a deep-sea setting characterized by thermogenic hydrocarbon seepage. Thermophilic endospores in permanently cold seabed sediments above petroleum seep conduits were correlated with the presence of hydrocarbons, revealing geofluid-facilitated cell migration pathways originating in deep oil reservoirs. Genomes of thermophilic bacteria highlight adaptations to life in anoxic petroleum systems and reveal that these dormant populations are closely related to oil reservoir microbiomes from around the world. After transport out of the subsurface and into the deep-sea, thermophilic endospores re-enter the geosphere by sedimentation. Viable thermophilic endospores spanning the top several metres of the seabed correspond with total endospore counts that are similar to or exceed the global average. Burial of dormant cells enables their environmental selection in sedimentary formations where new petroleum systems establish, completing a geological microbial loop that circulates living biomass in and out of the deep biosphere.

## Main text

Identifying natural forces that distribute organisms throughout the living world is critical to understanding Earth system functioning. Whereas the biogeography of animals and plants have been studied since the time of Darwin^5^, related ecological processes are harder to elucidate in the microbial realm where the effects of dispersal and environmental selection must be disentangled^6-8^. Dormant populations of microbes retain viability while enduring inhospitable conditions in relation to growth requirements, allowing dispersal to be studied directly without the influence of conflating factors like environmental selection. Bacterial endospores are equipped to survive dispersal over long distances and timescales^9^, with reports of viable spores ~2.5 km beneath the seafloor^10^ suggesting dispersal journeys lasting millions of years. This points to a genetically and functionally diverse seed bank of microbes that can be revived if subsurface environmental conditions select for their traits^11,12^.

The marine subsurface biosphere contains an estimated 10^29^ microbial cells contributing up to 2% of the total living biomass on Earth^1^. Whereas deep biosphere populations exhibit exponentially decreasing numbers with depth^13^, endospores experience less pronounced declines and appear to outnumber vegetative cells in deeper marine sediments^2,14^. Measurements of the endospore-specific biomarker dipicolinic acid indicate remarkably high numbers of endospores in deep warm strata, with depth profiles revealing that temperature influences sporulation and germination^13^. This is consistent with the prevalence of endospore forming *Firmicutes* in microbiome surveys of hot oil reservoirs from around the world^15^ where they actively contribute to biogeochemical cycling. In the energy limited deep biosphere, these petroleum systems represent energy rich oases^16,17^ that select for thermophilic organotrophy. Accordingly, cell densities in oil reservoirs can be an order of magnitude higher than those in surrounding sediments at the same depth^18^.

Hydrocarbon seepage up and out of deep petroleum systems is widespread in the ocean^19^. Studies of thermophilic spores in cold surface sediments globally^20,21^ have invoked warm-to-cold dispersal routes like hydrocarbon seeps to explain these observations^22^. In the Gulf of Mexico where cold seeps are common^23^, spore-forming thermophiles are correlated with the presence of migrated liquid hydrocarbons^24^ and buoyant gas migration mediates upward microbial dispersal in the top few centimeters^25^. Whether viable cells from deeper and hotter subsurface layers can be similarly circulated over greater depths and timescales by seepage and subsequent burial remains hypothetical. Here we compare deep-sea sediments from the NW Atlantic Ocean (Extended Data Fig. 1) using geophysics, hydrocarbon geochemistry, spore germination dynamics and genomics to demonstrate the dispersal cycle of viable cells throughout the marine subsurface. This geologically mediated microbial loop transports living biomass via upward seepage and downward burial and represents a previously overlooked mechanism for ecological maintenance and preservation of life in the energy limited subsurface biosphere.

Structural geology indicative of deep subsurface to surface geofluid conduits was determined by multichannel 2D and 3D seismic reflection surveys along the NW Atlantic Scotian Slope. Geophysical surveys covered ~70,000 km^2^ and obtained ~10,000 m of subsurface stratigraphic imagery in up to 3,400 m water depth (Fig. 1a). Seabed seep detection in deep-sea settings like this is very challenging^26^, thus a multi-disciplinary strategy was employed. Co-location at the seabed of the up-dip limit of deep-seated faults and seismic reflection anomalies considered to be direct hydrocarbon indicators were used to identify potential subsurface seep networks^27^ (Fig. 1b). These large-scale geophysical survey results were refined through high-resolution seismic reflection, side-scan sonar and multibeam bathymetry. Morphological features included a mounded structure with high backscatter intensity intersected by an elongated fracture-like depression (Fig. 1c) and a circular pockmark, suggesting the presence of a seep-like structure. Immediately beneath these features, high-resolution subsurface seismic profiling revealed a localized acoustic blanking zone (Fig. 1d) suggesting the presence of gas^28^ and a hydrocarbon migration pathway through subsurface sediments.

**Fig. 1.**
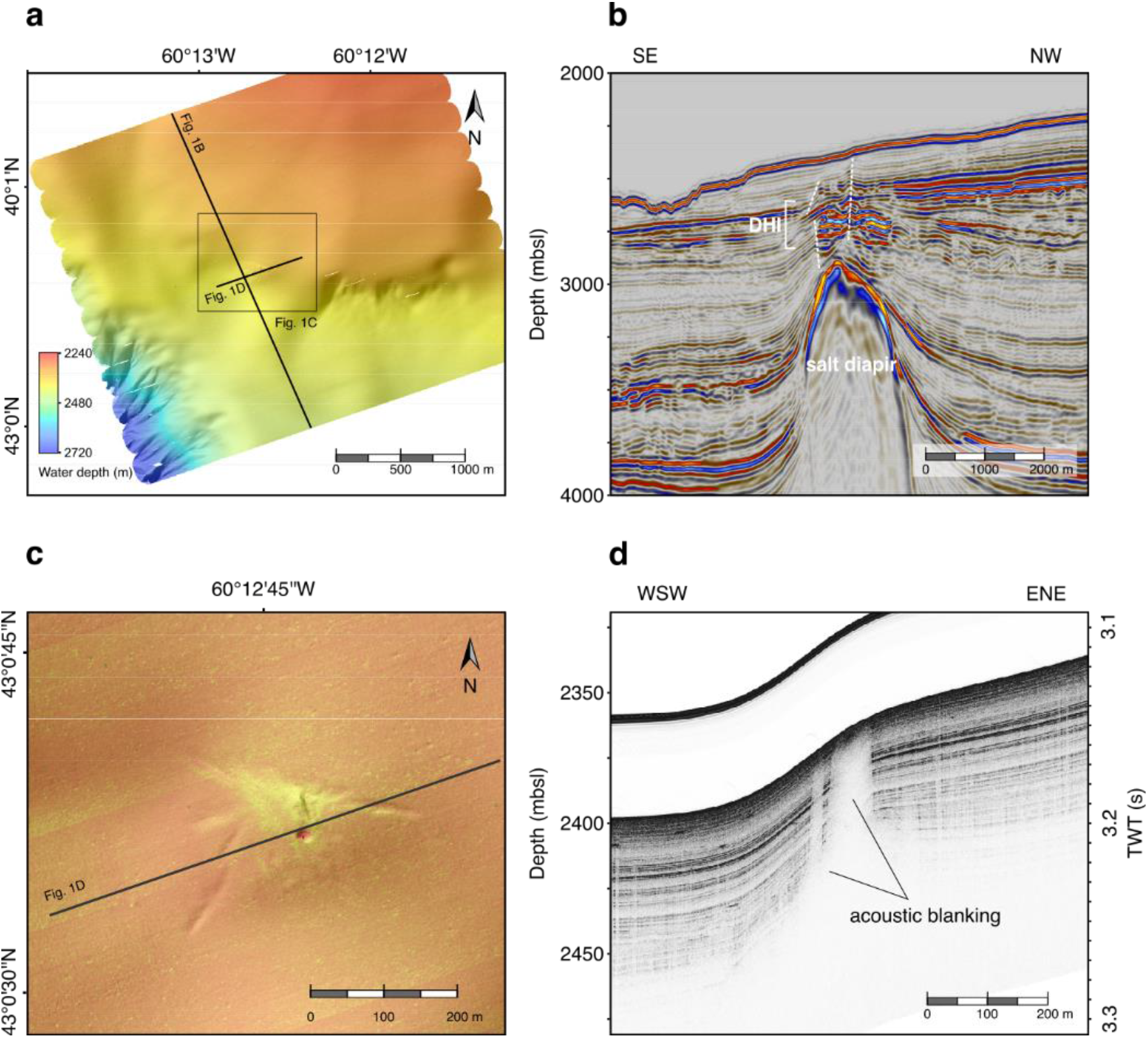
Deep subsurface to surface geofluid migration. **a,** Seafloor surface map derived from AUV-based multibeam bathymetric sonar data at one location. **b**, 3D seismic cross section showing a buried salt diapir, the location and direction of crestal faults (white dashed lines), and an interval with direct hydrocarbon indicators (DHI). **c**, Combined mosaic of side-scan sonar data and shaded relief bathymetry of the area surrounding a seep structure, indicating a pockmark feature as well as a small mound morphology. High backscatter intensity, related to distinctive properties of near-surface sediment, is shown in light-yellowish tones. **d**, AUV-based sub-bottom profiling showing localized acoustic blanking under the seep structure, indicative of upward fluid migration.

Locations showing seismic evidence of migrated hydrocarbons originating from a deep subsurface source (Supplementary Table 1) were examined in greater detail by comparing hydrocarbon signals from 14 different sites that were sampled by piston coring (Fig. 2a; Supplementary Table 2). Higher concentrations of thermogenic C_2_–*n*C_4_ compounds and heavy δ^13^C values for methane (−42 to −52‰) in interstitial gas, coupled with liquid hydrocarbon extracts featuring elevated *n*C_17_/*n*C_27_ ratios and a lack of odd-over-even alkane distributions in the *n*C_23-33_ range, provided clear evidence of migrated hydrocarbons at two sites. This was confirmed by higher proportions of thermally derived diasteranes relative to regular steranes (% 27 dβ S) and more thermally mature terpane distributions (C_30_ αβ relative to C_31_ αβ 22R hopane) in these cores. At the 12 other sites, thermogenic hydrocarbon signals were either inconclusive (*n*=4) or not detected (*n*=8).

**Fig. 2.**
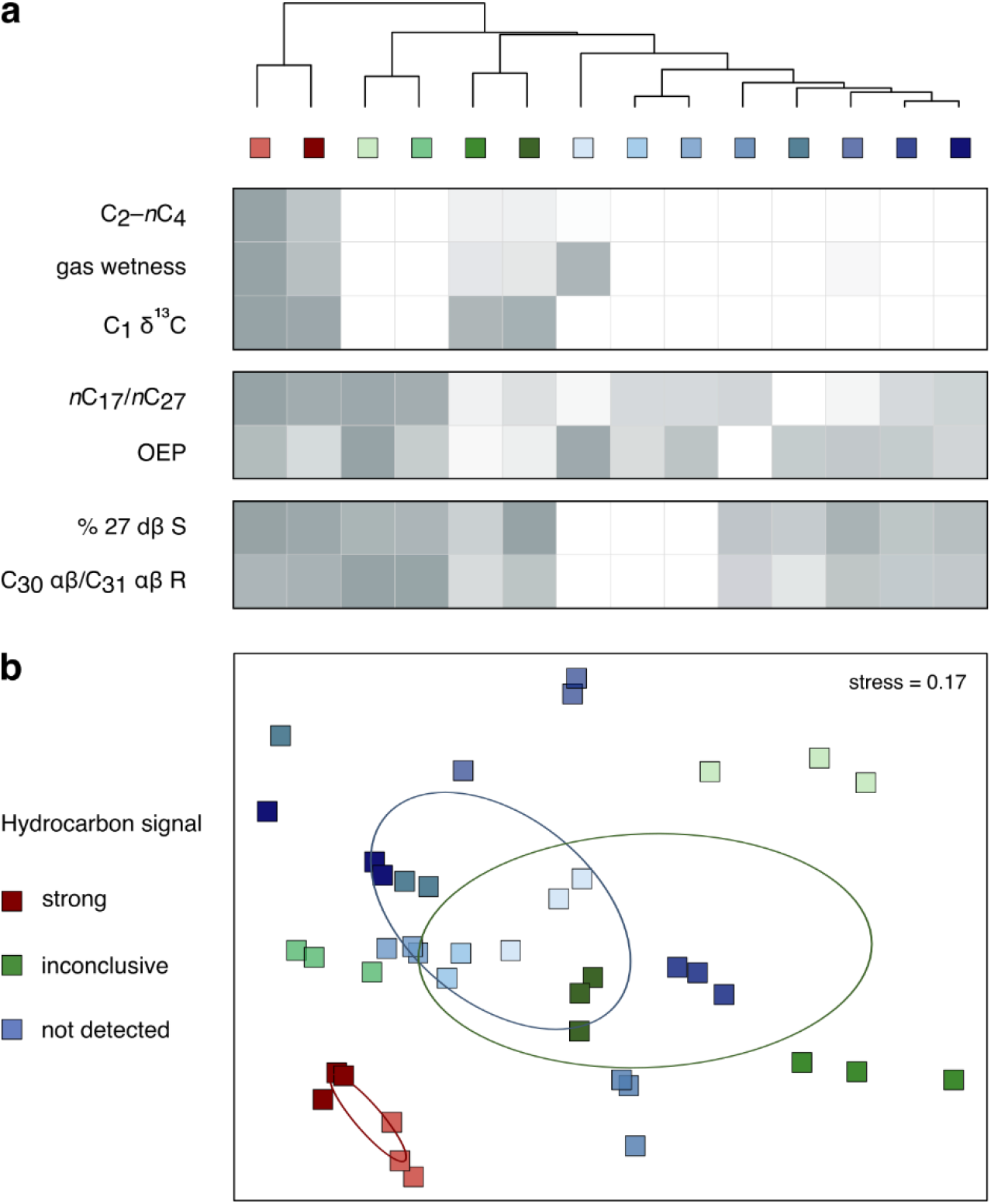
Hydrocarbon geochemistry and microbial community variance between seabed sampling sites. **a,** Gas (∑ C_2_–*n*C_4_, gas wetness, and C_1_ δ^13^C), liquid hydrocarbon extract (*n*C_17_/*n*C_27_ and odd-over-even predominance) and biomarker (% 27 dβ S and C_30_ αβ/C_31_ αβ 22R) measurements to assess the presence of thermogenic hydrocarbons. Each parameter is scaled between 0 and 1 as shown in the heatmap. Cores are represented by average values in instances where multiple depths from a core were tested (values provided in Supplementary Table 2). Hierarchical clustering highlights groups of sites where the evidence for the presence of thermogenic hydrocarbons is strong (red), inconclusive (green) or not detected (blue). **b**, Bray-Curtis dissimilarity in microbial community composition after sediment incubation at 50°C reflected the three geochemical groupings (ellipses indicate standard deviations of weighted averaged means of within-group distances for each of the three groups; see Extended Data Fig. 2 for plots from 40 and 60°C incubations). Sites with strong thermogenic hydrocarbon signals have distinct microbial populations after high temperature incubation (Supplementary Table 4).

To compare thermophilic spore-forming bacterial populations in cores with and without evidence of thermogenic hydrocarbons, endospore germination and thermophile enrichment was stimulated in high temperature anoxic incubations (40–60°C following pasteurization at 80°C; Extended Data Fig. 3). Assessing microbial community composition by 16S rRNA gene profiling of incubated surface sediments from all 14 locations showed divergent profiles for hydrocarbon-positive locations (Fig. 2b; Supplementary Table 3). Statistical comparisons revealed 42 unique amplicon sequence variants (ASVs), all belonging to spore-forming bacterial taxa, correlated with upward seepage of thermogenic hydrocarbons (IndicSpecies, P < 0.05; Supplementary Table 5). Putative fermentative organotrophs such as *Paramaledivibacter* and *Caminicella*, as well as sulfate-reducing *Desulfotomaculales* and *Candidatus* Desulforudis, showed strong hydrocarbon association (Fig. 3a). None of these groups were detected by applying the same DNA sequencing method to unincubated sediment (Supplementary Table 3), likely owing both to their low relative abundance *in situ*^29^ and their multi-layered endospore coat not yielding to standard cell lysis protocols for DNA extraction^30^.

**Fig. 3.**
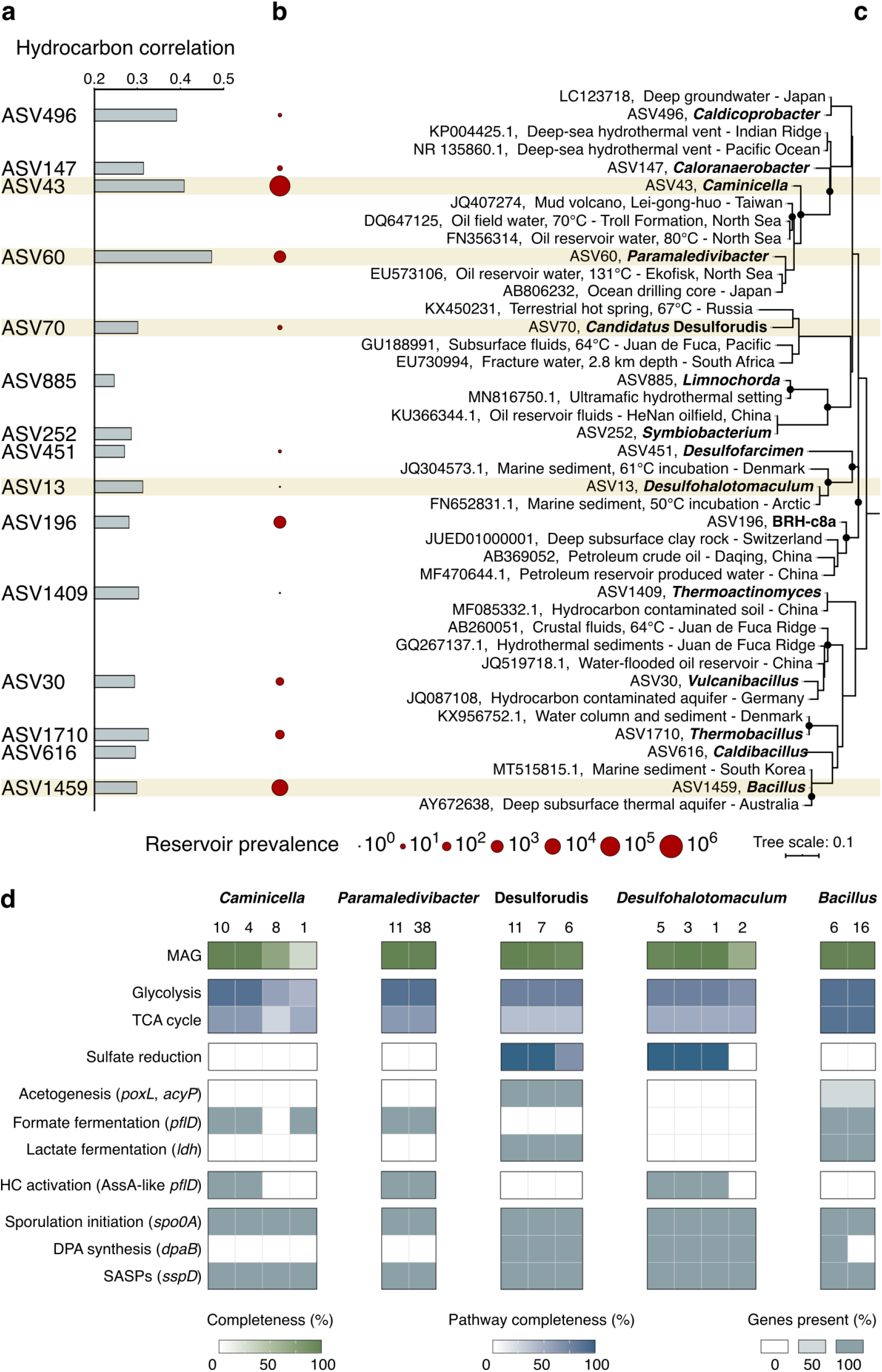
Oil reservoir provenance of seep-associated thermophiles. **a,** Correlation of thermophilic spore-forming bacterial amplicon sequence variants (ASVs) with thermogenic hydrocarbons. Highest ranking ASVs from each of 15 different genera are shown (representing 42 hydrocarbon-correlated ASVs in total). **b**, Prevalence of these genera in 59 oil reservoir microbiome assessments (11 million 16S rRNA gene sequences in total). **c**, Maximum likelihood phylogeny showing the 15 representative hydrocarbon-correlated ASVs and close relatives in the GenBank database (for all 42 indicator ASVs see Fig. S3). Black circles at the branch nodes indicate >80% bootstrap support (1,000 re-samplings), and the scale bar indicates 10% sequence divergence as inferred from PhyML. **d**, Metagenome-assembled genomes (MAGs) matching ASVs of interest (see corresponding brown shading in **a**–**c**) were assessed for anaerobic alkane degradation, sporulation and other metabolic features (see Supplementary Table 8 for pathway definitions).

To assess the prevalence of these bacteria in deep petroleum systems, we curated a dataset of 16S rRNA gene sequences from 59 different oil reservoir microbiomes from around the world (Supplementary Table 6). Seep-associated thermophilic endospore lineages identified in cold deep-sea sediments analysed here are found in high proportions in subsurface petroleum systems, especially *Caminicellaceae* and *Desulfotomaculales* which each make up 2–3% of the global oil reservoir microbiome dataset (Fig. 3b). ASV assessment at finer taxonomic resolution confirms close genetic relatedness between thermophilic endospores in Scotian Slope sediments and bacteria found in different subsurface oil reservoirs (Fig. 3c; Extended Data Fig. 4). Genomes of these dormant spores encode the potential for anaerobic hydrocarbon biodegradation, favouring their selection and growth in deep petroleum-bearing sediments (Fig. 3d). Metagenome-assembled genomes (MAGs) of *Caminicella, Paramaledivibacter, Desulfohalotomaculum* and *Bacillus* with rRNA sequences matching the indicator ASVs (Fig. 3; Supplementary Table 7) contain glycyl-radical enzymes proposed to mediate anaerobic alkane biodegradation via addition to fumarate^31,32^. Based on newly developed Hidden Markov Models for annotating alkylsuccinate synthases^33^, putative *assA* gene sequences in thermophilic spores diverge from canonical *assA* found in mesophilic *Proteobacteria* (Extended Data Fig. 5). This divergent clade includes thermophiles from hot oil reservoirs such as ^U^*Petromonas tenebris*^34^ and *Archaeoglobus fulgidus*^35^. Cold sediment MAGs also contain sporulation genes (Fig. 3d) including the *spo0A* master transcriptional response regulator^36^ as well as genes for synthesizing α/β-type small acid-soluble proteins (e.g., *sspD*) and dipicolinic acid (e.g., *dpaB*) involved in DNA protection^37^ (for a full list of sporulation genes see Supplementary Table 9).

Maintenance of dormancy has been proposed as a necessary pre-requisite for microbial taxa to exhibit biogeographic patterns over large distances and timescales^12^. Thermophilic endospores originating from deep petroleum-bearing sediments exemplify large-scale biogeography by connecting anaerobic hydrocarbon biodegradation and other microbial activities in the subsurface with intervening periods of large-scale migration in a dormant, sporulated state. Recurrent cyclical dispersal facilitates this scenario, consistent with a framework for microbial biogeography that features the same environment being both the origin and eventual destination for migrating populations^12^. Upon being transported out of the subsurface and into the benthos (Fig. 4a), further transport of spores via bottom water currents precedes eventual re-entry into the seabed (Fig. 4b). In the cold surface sediment of the Scotian Slope, thermophilic spores were detected in all of the cores that were collected, including those lacking geochemical evidence of hydrocarbon seepage (Supplementary Table 3). Dipicolinic acid concentrations within the top few metres demonstrate constant deposition and burial of endospores (Fig. 4c), with numbers in this region similar to or exceeding the seabed global average^2^. High temperature anoxic incubation of sediment from these depths to germinate thermophilic spores shows that they remain viable during burial (Extended Data Fig. 6). In sediments that eventually become petroleum systems at even greater depths where these temperatures occur naturally, the activation of these dormant bacteria by suitable nutrients and heat completes a subsurface microbial loop of viable cells circulating out of and back into the deep biosphere (Fig. 4).

**Fig. 4.**
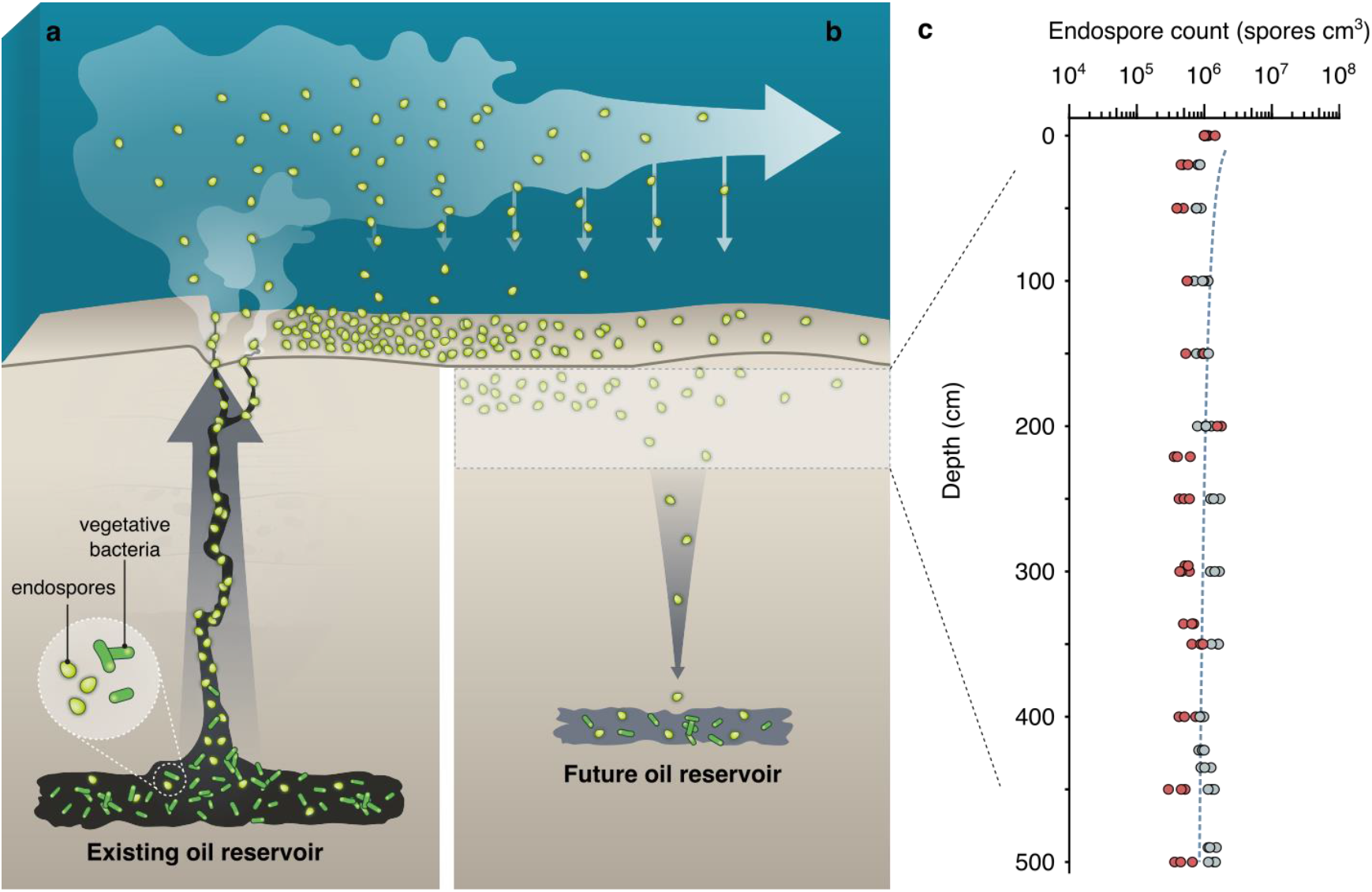
Subsurface microbial loop mediated by seepage, sedimentation, dormancy and environmental selection. **a,** Endospore-forming microbial populations actively inhabiting deep petroleum systems get dispersed upwards as dormant spores by hydrocarbon seepage along geological conduits. Endospores entering the deep-sea are dispersed laterally by bottom water currents. **b**, Endospores get deposited on the seabed and undergo burial. **c**, Endospore burial is revealed by triplicate measurements of dipicolinic acid concentrations in the upper few metres of the seabed in two Scotian Slope sediment cores where hydrocarbons were not detected. The dashed regression line reflects the global average estimated for endospores in the marine subseafloor biosphere^2^. Survival of some fraction of these endospores over long time-scales^10,38^ enables environmental selection (i.e., germination and activity) by suitable substrates and heat—favorable conditions that establish in sediments where oil migrates into to establish a reservoir^39^. The sequence shown in **a** and **b** completes a ‘subsurface microbial loop’ that incorporates cell dispersal and biogeochemical cycling in Earth’s deep biosphere.

Subsurface marine sediments contain 12–45% of Earth’s microbial biomass and are central to the planet’s biogeochemical cycling^40^. This is especially true in petroleum systems that control and are controlled by subsurface microbial populations^31,41^. Despite the importance of these processes, research on the subsurface microbiome rarely focuses on ecological factors like dispersal and selection, preventing a more complete understanding of deep biosphere ecosystems^4^. The results presented here demonstrate that geological processes of geofluid flow and sedimentation connect deep petroleum systems with the ocean and mediate a recurrent and spatially extensive cycle of microbial dispersal throughout the subsurface. This circulation of living biomass is uniquely characterized by defined episodes of microbial activity in petroleum-bearing sediments interspersed by long intervals of passive dispersal — an ecological sequence that is difficult to delineate as clearly in other environmental settings^8^. By connecting the physical and physiological factors that govern survival and evolution in the deep biosphere, this subsurface microbial loop showcases the geosphere as a model system for understanding the interplay between microbial dispersal and selection in the biosphere at large spatial and temporal scales.

## Methods

### Seismic data acquisition and processing

Multiple two- and three-dimensional multi-channel seismic surveys performed here for the identification of seafloor seeps relied on an earlier regional 28,000 km^2^ 2D seismic survey. The earlier survey was shot in a 6 km grid, acquiring 14 seconds of data with 80–106-fold and a 2 millisecond sampling interval. The 1998 vintage used was processed to pre-stack time migrated data. 2D seismic survey interpretations were refined using the Shelburne 3D Wide Azimuth Seismic survey. This survey was acquired over 12,000 km^2^ in the deep-water Shelburne sub-basin at 6.25 x 50 m bin spacing with a fold of 100. This vintage of data utilized both full 3D Anisotropic Kirchhoff pre-stack time migration (PSTM) and full volume anisotropic Kirchhoff pre-stack depth migration (PSDM) with vertical transverse isotropy. PSTM had a processed bin size of 12.5 x 25 m, while PSDM had an output bin size of 25 x 25 m. Data were interpreted using the Petrel E&P Software Platform (Schlumberger Limited).

High-resolution seismic reflection profiles (data not shown) were used to investigate the subsurface stratigraphy in the vicinity of seep prospects to inform autonomous underwater vehicle (AUV) survey and coring locations. Profiles were collected during three expeditions between 2015 and 2018 onboard the CCGS *Hudson*^42–44^ using a Huntec single-channel Deep Tow Seismic (DTS) sparker system. Tow depth was ~100 m beneath the sea surface with the source fired at a moving time interval between 1 and 3 seconds. The peak frequency for the Huntec DTS sparker is approximately 1,500 Hz and spans from 500–2,500 Hz. Raw sparker data was processed using the VISTA Desktop Seismic Data Processing Software (Schlumberger Limited) and included Ormsby band-pass filtering, scaling correction, automatic gain control and trace mixing.

An autonomous underwater vehicle (AUV) was deployed from the vessel *Pacific Constructor* to collect high-resolution geophysical data over a 2.5 x 2.5 km area at the location of cores 16-41 and 18-7 in August 2020. The HUGIN 6000 AUV (Kongsberg Maritime) was steered approximately 40 m above the seafloor for multibeam bathymetric, side-scan sonar and sub-bottom profiling data collection using a Kongsberg EM 2040 multibeam echosounder and a EdgeTech 2205 sonar system, respectively. EM2040 Multibeam bathymetric data was acquired at a frequency of 400 kHz, with a continuous waveform (CW) pulse and synchronized with Doppler velocity log. Multibeam bathymetric data was processed using Caris and Eiva suite. Side Scan Sonar data was acquired at a frequency of 230 kHz and post-processing of the data was completed in Sonarwiz. The sub-bottom profiler was operated over the frequency range of 1–9 kHz with a 20-millisecond pulse. The high-resolution seismic data was integrated and analyzed using IHS Kingdom Suite (IHS Markit Ltd). Acoustic travel times for high resolution sub-bottom profiler lines were converted into depths by using an average seismic velocity of 1,500 m.s^-1^.

### Marine sediment sampling

Seabed surface sediments in 2,000 to 3,400 m water depth were collected by piston and gravity coring from different locations on the Scotian Slope, offshore Nova Scotia, Canada (Fig. 1; Supplementary Table 1) during May-June expeditions aboard the CCGS *Hudson* in 2015, 2016 and 2018^42-44^. Piston cores, trigger weight cores (smaller cores that release the head weight of the piston core) and gravity cores ranged from 0.18 to 8.34 m in length. Upon recovery, cores were split longitudinally onboard the ship. Sediment intervals from the base of the core (5-10 cm) were transferred to gas-tight IsoJars^®^ (Isotech Laboratories Inc., USA), immediately flushed with nitrogen, and stored at −20°C prior to interstitial gas analysis. Similar intervals from variable depths along the cores, selected based on indications of visible hydrocarbon staining or odour, fluorescence, or sandy lithology, were stored in aluminum foil at −20°C for eventual hydrocarbon analysis, and in sterile Whirl-Pak^®^ bags or glass jars at 4°C for eventual high temperature endospore germination experiments. Sediment intervals at the top of the core (either 0–10 or 0–20 cm below seabed) were similarly transferred to sterile Whirl-Pak^®^ bags or sterile glass jars.

### Hydrocarbon geochemical analysis

Interstitial gas analysis was performed on aliquots of IsoJar^®^ (Isotech Laboratories Inc., USA) headspace transferred into Exetainers^®^ (Labco Limited, UK). Sample volumes of 1 mL were injected into an Agilent 7890 RGA Gas Chromatograph (Agilent Technologies, USA). A flame ionisation detector determined C_1_–C_5_ hydrocarbon gas concentrations that were used to calculate gas wetness. Carbon isotopic composition (δ^13^C) of hydrocarbon gas components was determined by gas chromatography combustion isotope ratio mass spectrometry; headspace aliquots were analyzed on a Trace 1310 Gas Chromatograph (Thermo Fisher Scientific, USA) interfaced to a Delta V Isotope Ratio Mass Spectrometer (Thermo Fisher Scientific, USA).

Sediments were analyzed for hydrocarbon biomarkers in subsamples where sufficient extract yields were recovered. Accordingly, no extract yield, or insufficient yields to determine biomarker concentrations, were considered indicative of the absence of hydrocarbon seepage. Organic matter was extracted from sediment by adding dichloromethane with 7% (v/v) methanol, mixing the solution in an ultrasonic bath for 15 min and then leaving at room temperature for 24 h. Extractable organic matter (EOM) was evaporated to dryness and weighed. Asphaltenes were removed by pentane addition in excess (40 times the volume of EOM), storage for 12 h, and centrifugation. Gas chromatography analysis of the EOM was performed on an Agilent 7890A Gas Chromatograph (Agilent Technologies, USA). Saturate and aromatic hydrocarbon fractions showing possible evidence of thermogenic hydrocarbons were analyzed further using a Micromass ProSpec Gas Chromatography-Mass Spectrometer (Waters Corporation, USA). Geochemical analyses were performed by Applied Petroleum Technology, Norway, to the standards used in industrial hydrocarbon assessments. Geochemistry data was collectively interpreted for evidence of thermogenic hydrocarbons likely derived from subsurface hydrocarbon seeps. Hierarchical clustering (complete linkage clustering based on Euclidean distance) of geochemical measurements scaled between 0 and 1 over the range of values (0 representing weakest thermogenic signal and 1 representing the strongest thermogenic signal) was used to further assess and visualise groups of sites with similar geochemical signatures (Fig. 2a).

### Sediment incubation at elevated temperatures

Sediments were investigated for the germination and growth of dormant bacterial endospores. Following homogenizing by stirring within the sample container, up to 100 g of sediment was transferred into separate 250 mL serum bottles that were sealed with butyl rubber stoppers (Chemglass Life Sciences, Canada) and the headspace exchanged with N_2_:CO_2_ (90:10%). Sediment slurries were prepared in a 1:2 (w/w) ratio with sterile, anoxic, synthetic seawater medium^45^ containing 20 mM sulfate and amended with acetate, butyrate, formate, lactate, propionate, and succinate (5 mM each for surface sediments and 1 mM each for deeper sediments). Master sediment slurries were subdivided into replicate, sterile, anoxic 50 mL serum bottles sealed with butyl rubber stoppers. Slurries were pasteurized at 80°C for 1.5 h to kill vegetative cells and select for heat-resistant endospores. Triplicate pasteurized slurries were immediately incubated at 40, 50 or 60°C for up to 56 days to promote germination and growth of thermophilic endospore-forming bacteria. Subsamples (2 mL) were periodically removed using sterile N_2_:CO_2_-flushed syringes and stored at −20°C for molecular analysis.

### 16S rRNA gene amplicon sequencing

Genomic DNA was extracted from triplicate slurries subsampled immediately before incubation (i.e., post-pasteurization), and periodically during the incubation, using the DNeasy PowerLyzer PowerSoil Kit (Qiagen, USA). Extractions were performed on 300 μL of slurry according to the manufacturer’s protocol, except for inclusion of a 10 min incubation at 70°C immediately after the addition of Solution C1 to enhance cell lysis. Extraction blanks (Milli-Q water) were processed in parallel. DNA was quantified using the Qubit dsDNA High Sensitivity assay kit on a Qubit 2.0 fluorometer (Thermo Fisher Scientific, Canada). The V3 and V4 hypervariable regions of the 16S rRNA gene were amplified in triplicate PCR reactions per extraction using the primer pair SD-Bact-341-bS17/SD-Bact-785-aA21^46^ modified with Illumina MiSeq overhang adapters. All PCR reactions were performed in triplicate. All DNA extraction blanks and PCR reagent blanks were confirmed for negative amplification using agrose gel electrophoresis. Triplicate PCR products were pooled, purified using a NucleoMag NGS Clean-up and Size Select kit (Macherey-Nagel Inc., USA) and indexed. Sizes of indexed amplicons were verified using the High Sensitivity DNA kit on an Agilent 2100 Bioanalyzer system (Agilent Technologies, Canada). Indexed amplicons were pooled in equimolar amounts and sequenced on an in-house Illumina MiSeq benchtop sequencer (Illumina Inc., USA) using Illumina’s v3 600-cycle reagent kit to obtain 300 bp paired-end reads.

### 16S rRNA gene amplicon sequence processing

A total of 20,589,990 raw paired-end reads were generated across six separate MiSeq runs. Primers were trimmed using Cutadapt version 2.7^47^ prior to amplicon sequence variant (ASV) inference using DADA2 version 1.16^48^ in base R version 3.6.1^49^. Forward and reverse read pairs were trimmed to a run-specific length defined by a minimum quality score of 25. Read pairs were filtered allowing no ambiguous bases and requiring each read to have less than two expected errors, and PhiX sequences removed. Reads were dereplicated providing unique sequences with their corresponding abundance. Error rates were estimated from sequence composition and quality by applying a core denoising algorithm for each sequencing run to account for run-to-run variability. Unique ASVs were inferred independently from the forward and reverse reads of each sample, using the run-specific error rates, and then pairs were merged if they overlapped with no mismatches. Chimeras were identified and removed, then an additional length trimming step removed sequence variants shorter than 400 nucleotides and larger than 435 nucleotides. A total of 32,018 ASVs were resolved from 11,355,683 quality-controlled reads. Taxonomy was assigned using the Ribosomal Database Project’s k-mer-based naïve Bayesian classifier with the DADA2-formatted Silva database version 138^50^. Reads were randomly subsampled without replacement to the smallest library size (*n*=4,635) using the *phyloseq* R package^51^ prior to comparative analysis.

### Metagenome sequencing

Genomic DNA extracted from four separate sediment slurries after 56 days of incubation was used for metagenomic sequencing. Library preparation and sequencing was conducted at the Center for Health Genomics and Informatics in the Cumming School of Medicine, University of Calgary. DNA was sheared using a Covaris S2 ultrasonicator (Covaris, USA), and fragment libraries prepared using a NEBNext Ultra II DNA Library Prep Kit for Illumina (New England BioLabs, USA). Metagenomic libraries were sequenced on the Illumina NovaSeq platform (Illumina Inc., USA) using an S4 flow cell with Illumina 300 cycle (2 × 150 bp) V1.5 sequencing kit.

### Metagenome sequence processing

A total of 65,786,766 raw reads from four metagenomic libraries were quality-controlled by trimming technical sequences (primers and adapters) and low-quality additional bases, and filtering artifacts (phiX), low-quality reads and contaminated reads using BBDuk (BBTools suite, http://jgi.doe.gov/data-and-tools/bbtools). Trimmed and filtered reads from each metagenome were assembled separately, as well as co-assembled, using MEGAHIT version 1.2.2^52^ using default parameters and with <500 bp contigs removed. Binning of the four assemblies and one co-assembly was performed using MetaBAT 2 version 2.12.1^53^. Contamination and completeness of the resulting MAGs were estimated using CheckM version 1.0.11^54^ with the lineage-specific workflow. Ribosomal rRNA genes were identified in unbinned reads using phyloFlash^55^ and in binned reads using rRNAFinder implemented in MetaErg version 1.2.0^56^. Protein coding genes were predicted and annotated against curated protein sequence databases (Pfam, TIGRFAM, and Swiss-Prot) using MetaErg version 1.2.0^56^. Metabolic pathways were identified using KEGG Decoder^57^ to parse genes annotated with KEGG Orthology using BlastKOALA^58^. Hydrocarbon degradation genes were additionally annotated using CANT-HYD^33^ following gene predictions made using Prodigal version 2.6.3^59^. MAGs were classified with GTDB-Tk version 1.3.0^60^ and by alignment with Silva database version 138^50^ using mothur version 1.39.5^61^ in instances where 16S rRNA gene was recovered by rRNAFinder^56^.

MAGs for the seep-associated taxa were identified by alignment of predicted 16S rRNA gene sequences recovered from bins with the seep indicator ASV sequences highlighted by IndicSpecies (see below). An alignment identity of 100% across the full length of the amplicon was required to confirm association. In instances where a V3-V4 overlapping 16S rRNA gene sequence was not recovered in the MAG, taxonomic classification of the partial 16S rRNA gene, the MAG (GTDB-Tk version 1.3.0^60^), or the sample by phyloFlash was used to identify possible associations to seep-associated taxa. If the most abundant ASV in an unrarefied 16S rRNA gene amplicon library, with the same taxonomic classification as the recovered 16S rRNA gene or MAG, corresponded to the most abundant ASV in that sample, a probabilistic association was assumed and the MAG was retained for further analysis. Replicate MAGs were identified from cluster groups based on metagenome distance estimation using a rapid primary algorithm (Mash) and average nucleotide identity (ANI) using dRep version 2.3.2^62^ and included in the analysis.

### Analysis of global oil reservoir microbiome sequences

Raw high-throughput sequence data, totalling 53,019,792 reads from ten separate studies, was obtained from the National Center for Biotechnology Information’s (NCBI) Sequence Read Archive^63^ (SRA) by compiling sequence accession lists and using the SRA Toolkit. Initial sequence data processing was performed using VSEARCH version 2.11.1^64^. If necessary, paired-end sequence files were merged based on a minimum overlap length of 10 base pairs (bp) and a maximum permitted mismatch of 20% of the length of the overlap. Merged reads were filtered with a maximum expected error of 0.5 for all bases in the read, and minimum and maximum read lengths of 150 and 500 bp, respectively. Identical reads were dereplicated and annotated with their associated total abundance for each sample, prior to *de novo* chimera detection. Re-replication resulted in 10,857,433 quality-controlled reads. In addition to these amplicons generated by high-throughput sequencing platforms, 2,850 near full length amplicon sequences from 49 separate clone library and/or cultivation-based studies were downloaded from the NCBI’s GenBank database using published accession numbers. Taxonomy was assigned to the combined 10,860,283 sequences using the Ribosomal Database Project’s k-mer-based naïve Bayesian classifier with the Silva database version 138^50^.

### Statistical analysis and data visualization

Statistical analyses and visualization were performed using base R version 3.6.1^49^, or the specific R packages described below. Non-metric multidimensional scaling (NMDS) of Bray-Curtis dissimilarity was calculated using the *metaMDS* function of the *vegan* package^65^ in R and visualized using the *ggplot2* package^66^. Analysis of similarity (ANOSIM) tests measured significant differences between sediment communities and were performed using the *anosim* function of the *vegan* package^65^.

Microbial indicator sequence analysis, designed to test the association of a single ASV with an environment through multilevel pattern analysis, was used to identify sequences that best represent specific sediments or groups of sediments under variable test conditions based on both ASV presence/absence and relative abundance patterns. Indicator ASVs were calculated using the *multipatt* function of the *indicspecies* package in R, employing a point-biserial correlation index ^67^. Tests were performed on amplicon libraries constructed after 28 and 56 days of high temperature incubation, omitting pre-incubation (day-0) libraries as representing samples prior to endospore enrichment. Among the 32,018 ASVs, only those present in >1% relative abundance in at least one sample across the entire dataset were included in the analysis. The strength of the association is represented by the IndicSpecies Stat value (plotted in Fig. 3a). Only observations with *P* < 0.05 were considered statistically significant and reported.

### Phylogenetic analysis

ASVs associated with thermogenic hydrocarbons, together with their five most closely related sequences from Genbank (determined by BLAST searches), were aligned using the web-based multiple sequence aligner SINA^68^. Aligned sequences were imported into the ARB-SILVA 138 SSU Ref NR 99 database^50^ and visualized using the open-source ARB software package^69^. A maximum likelihood (PhyML) tree was calculated with near full length (>1,300 bases) bacterial 16S rRNA gene reference sequences as well as those from closest cultured isolates. In total, 172 sequences were used to calculate phylogeny (bootstrapped with 1,000 re-samplings), accounting for 1,006 alignment positions specified based on positional variability and termini filters for bacteria. Using the ARB Parsimony tool, ASV and Genbank sequences were added to the newly calculated tree using positional variability filters covering the length of the representative sequences for each sequence without changing the overall tree topology (Extended Data Fig. 4). Trees were annotated using iTOL version 5.5^70^.

### Dipicolinic acid (DPA) measurement

Sediment samples were prepared in triplicate using the methods described in Lomstein and Jørgensen (2012)^71^ and Rattray *et al.* (2021)^72^. To extract DPA, 0.1 g of freeze-dried sediment was hydrolysed by addition of 6M HCl and heating at 95°C for 4 hours, before quenching on ice to stop hydrolysis. The hydrolysate was freeze dried, reconstituted in Milli-Q water, frozen and freeze dried again. Samples were then dissolved in 1M sodium acetate and aluminium chloride was added. Sediment extracts were filtered (0.2 μm) and mixed with terbium (Tb^3+^) prepared in 1M sodium acetate. DPA was separated and eluted using gradient chromatography over a Kinetex 2.6 μm EVO C18 100Å LC column (150 x 4.5 mm; Phenomenex, USA) fitted with a guard column. Solvent A was 1M sodium acetate amended with 1M acetic acid to pH 5.6 and solvent B was 80% methanol: 20% water pumped with a Thermo RS3000 pump (Thermo Scientific Dionex, USA). The sample injection volume was 50 μl and the total run time was 10 min (including flushing). Detection was performed using a Thermo FLD-3000RS fluorescence detector (Thermo Scientific Dionex, USA) set at excitation wavelength 270 nm and emission 545 nm. To determine DPA concentrations under the limit of detection, samples were analysed using standard addition^71^. For this, a known concentration of DPA standard /Tb^3+^ sodium acetate was sequentially added to the sediment exact and analysed. Concentrations were calculated using methods described in Lomstein and Jørgensen (2012)^71^.

#### Data and materials availability

All data is available in the main text or the supplementary materials. Amplicon and metagenome sequences generated in this study are available through the NCBI Sequence Read Archive (https://www.ncbi.nlm.nih.gov; BioProject accession number PRJNA604781).

## Acknowledgements

We wish to thank the crew of CCGS *Hudson* and Natural Resources Canada for collection of piston cores and onboard core processing. Ship time funding and support was provided by the Nova Scotia Department of Energy and Mines, the Nova Scotia Offshore Energy Research Association, and Natural Resources Canada. The authors wish to thank Steve Larter, Lisa Gieg, Bo Barker Jørgensen and Rhonda Clark for helpful discussions and suggestions and research support.

## Author contributions

C.R.J.H. and A.M. secured research funding. P-A.D, N.M., D.C.C. and A.M. acquired, processed and analyzed geophysical data. D.A.G., A.C., C.L., M.A.C., J.W., A.M., D.C.C. and C.R.J.H. collected and processed marine sediment samples for testing. M.F., J.W. and D.A.G. generated and analyzed hydrocarbon geochemical data. D.A.G., A.C., C.L., O.H., M.A.C. and C.R.J.H generated, processed and interpreted microbial community data. D.A.G. and S.B. curated and analyzed reservoir microbiome data. D.A.G. and J.Z. processed and analyzed metagenomic data. J. R. measured and interpreted DPA signals. D.A.G. and C.R.J.H. drafted the manuscript with feedback from all authors during refinement and finalization.

## Additional information

### Correspondence and requests for materials

can be addressed to D.A.G. and C.R.J.H.

