## Extended Data for "Geological processes mediate a subsurface microbial loop in the deep biosphere"

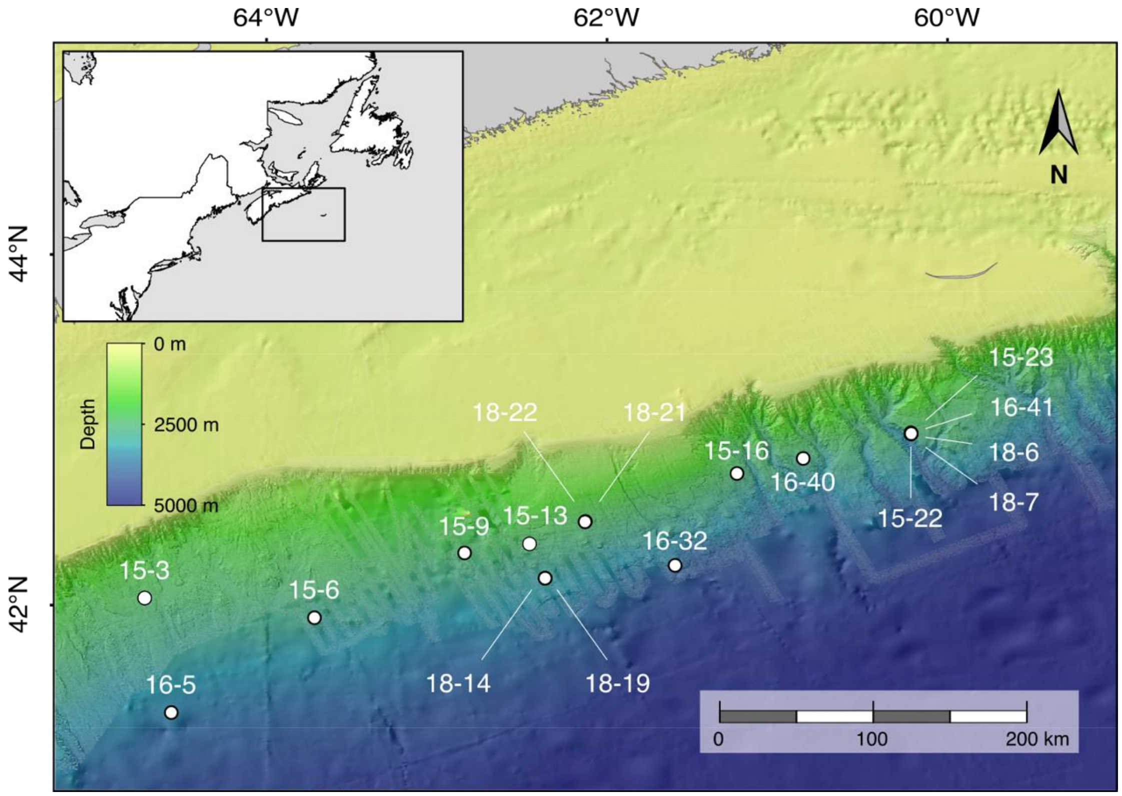


**Extended Data Fig. 1 | Deep sea study sites in the NW Atlantic Ocean.** Sediment coring locations on the Scotian Slope. The inset shows the extent of the 20,000 km$^2$ study area, off the east coast of Nova Scotia, Canada. Bathymetric map from the General Bathymetric Chart of the Oceans (GEBCO, www.gebco.net) and National Oceanic and Atmospheric Administration (NOAA, www.ngdc.noaa.gov/).

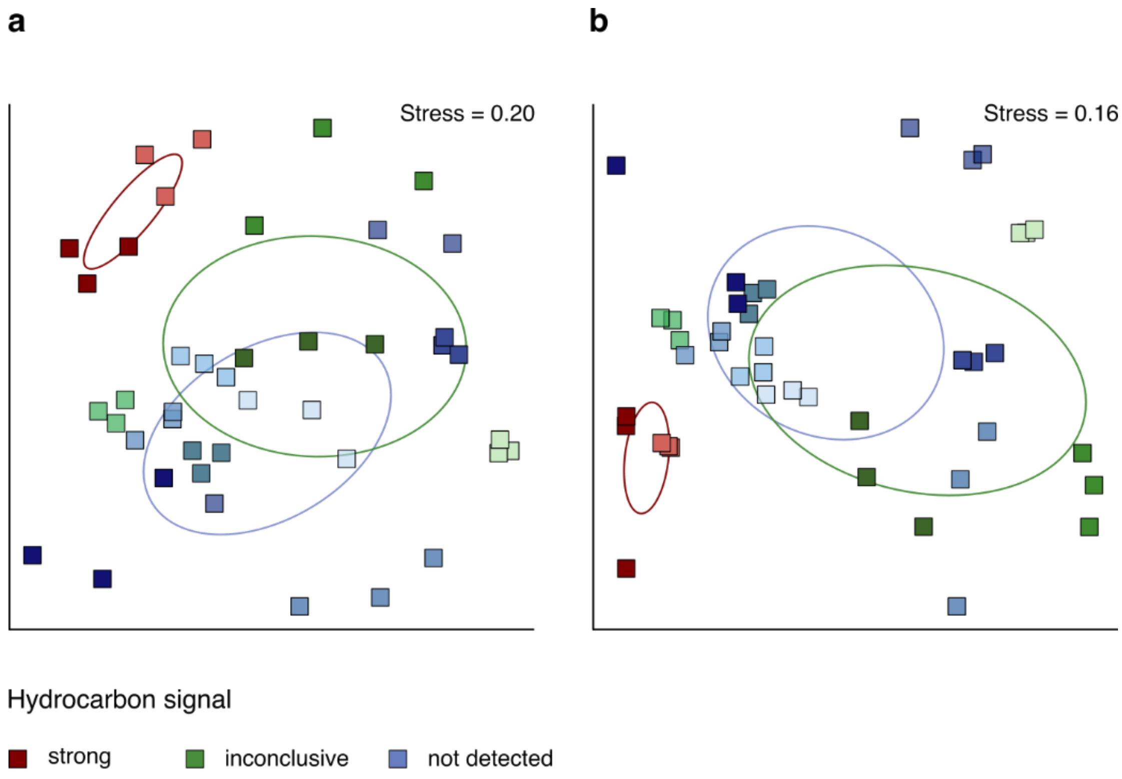


**Extended Data Fig. 2 | Microbial community variance between core sites.** Non-metric multidimensional scaling of the Bray-Curtis dissimilarity of the microbial community composition after sediment incubation at (**a**) 40°C and (**b**) 60°C (for 50°C incubations, see Fig. 2b). Red symbols indicate sites with strong geochemical evidence of hydrocarbons (*n*=2), green symbols indicate sites with inconclusive hydrocarbon signals (*n*=4), and blue symbols indicate sites where hydrocarbons were not detected (*n*=8). Triplicate amplicon libraries are plotted for each condition. Oil-positive locations have distinct microbial populations as indicated by standard deviation ellipses of the hydrocarbon groups.

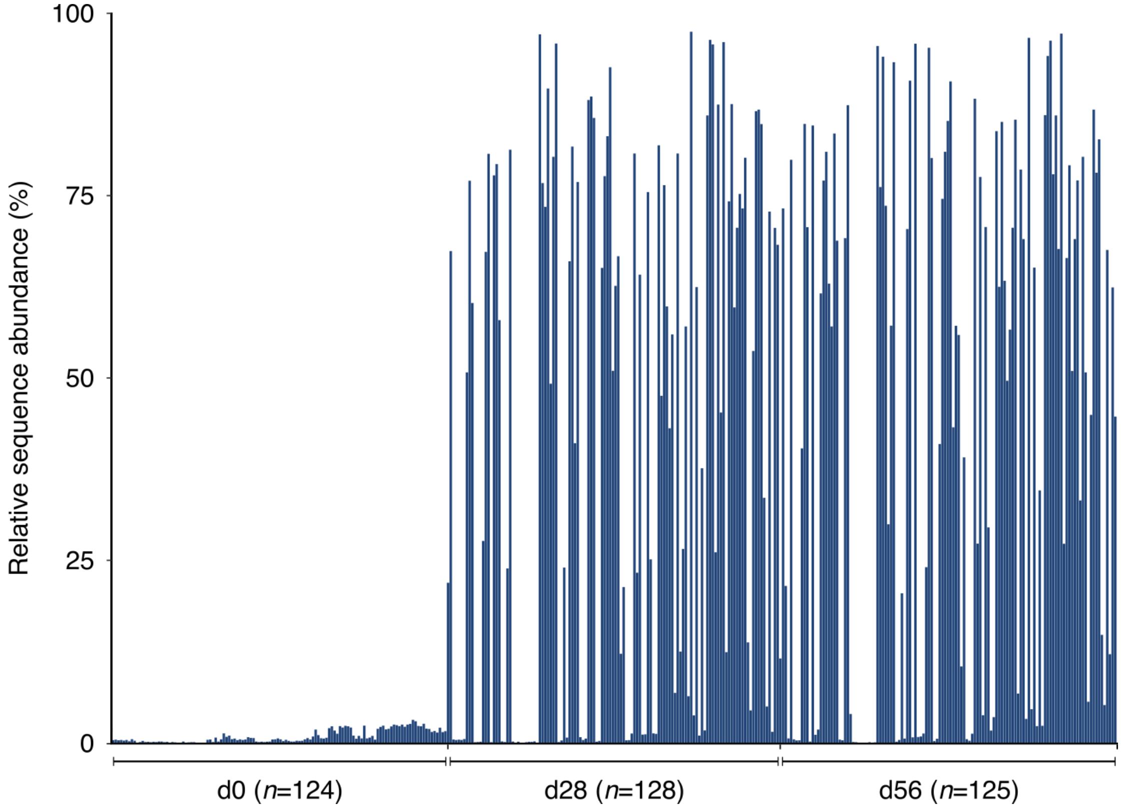


**Extended Data Fig. 3 | Endospore germination and enrichment.** Relative sequence abundance of *Firmicutes* in 372 rRNA gene amplicon libraries before (d0) and after (d28 and d56) high temperature anoxic incubation. Each blue bar represents a single library of 4,635 subsampled reads and demonstrates an increased proportion of *Firmicutes* following 28- and 56-days of incubation.

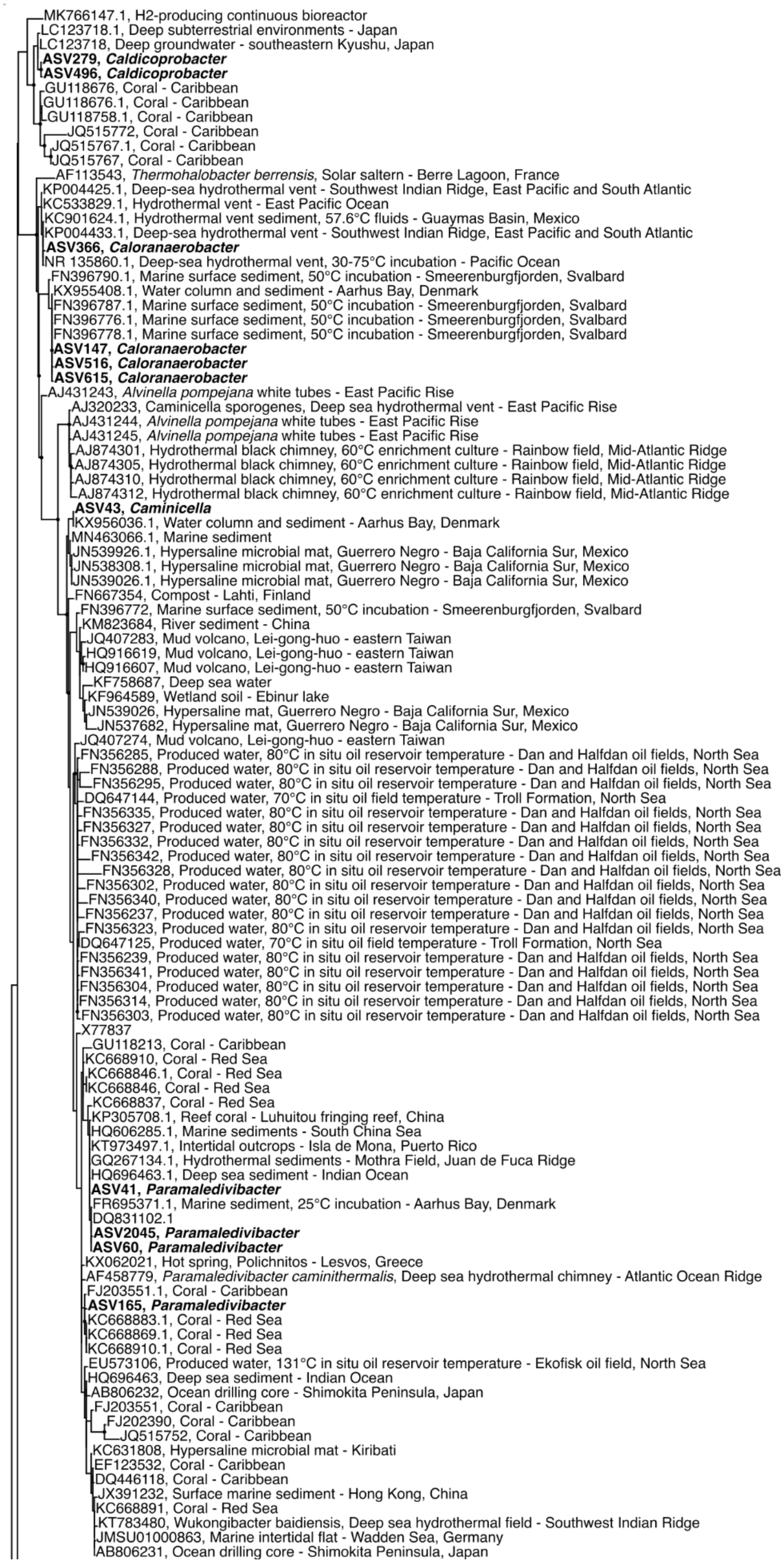
MK766147.1, H2-producing continuous bioreactor
LC123718.1, Deep subterrestrial environments - Japan
LC123718, Deep groundwater - southeastern Kyushu, Japan
**ASV279, *Caldicoprobacter***
**ASV496, *Caldicoprobacter***
GU118676, Coral - Caribbean
GU118676.1, Coral - Caribbean
GU118758.1, Coral - Caribbean
JQ515772, Coral - Caribbean
JQ515767.1, Coral - Caribbean
JQ515767, Coral - Caribbean
AF113543, *Thermohalobacter berrensis*, Solar saltern - Berre Lagoon, France
KP004425.1, Deep-sea hydrothermal vent - Southwest Indian Ridge, East Pacific and South Atlantic
KC533829.1, Hydrothermal vent - East Pacific Ocean
KC901624.1, Hydrothermal vent sediment, 57.6°C fluids - Guaymas Basin, Mexico
KP004433.1, Deep-sea hydrothermal vent - Southwest Indian Ridge, East Pacific and South Atlantic
**ASV366, *Caloranaerobacter***
NR 135860.1, Deep-sea hydrothermal vent, 30-75°C incubation - Pacific Ocean
FN396790.1, Marine surface sediment, 50°C incubation - Smeerenburgfjorden, Svalbard
KX955408.1, Water column and sediment - Aarhus Bay, Denmark
FN396787.1, Marine surface sediment, 50°C incubation - Smeerenburgfjorden, Svalbard
FN396776.1, Marine surface sediment, 50°C incubation - Smeerenburgfjorden, Svalbard
FN396778.1, Marine surface sediment, 50°C incubation - Smeerenburgfjorden, Svalbard
**ASV147, *Caloranaerobacter***
**ASV516, *Caloranaerobacter***
**ASV615, *Caloranaerobacter***
AJ431243, *Alvinella pompejana* white tubes - East Pacific Rise
AJ320233, Caminicella sporogenes, Deep sea hydrothermal vent - East Pacific Rise
AJ431244, *Alvinella pompejana* white tubes - East Pacific Rise
AJ431245, *Alvinella pompejana* white tubes - East Pacific Rise
AJ874301, Hydrothermal black chimney, 60°C enrichment culture - Rainbow field, Mid-Atlantic Ridge
AJ874305, Hydrothermal black chimney, 60°C enrichment culture - Rainbow field, Mid-Atlantic Ridge
AJ874310, Hydrothermal black chimney, 60°C enrichment culture - Rainbow field, Mid-Atlantic Ridge
AJ874312, Hydrothermal black chimney, 60°C enrichment culture - Rainbow field, Mid-Atlantic Ridge
**ASV43, *Caminicella***
KX956036.1, Water column and sediment - Aarhus Bay, Denmark
MN463066.1, Marine sediment
JN539926.1, Hypersaline microbial mat, Guerrero Negro - Baja California Sur, Mexico
JN538308.1, Hypersaline microbial mat, Guerrero Negro - Baja California Sur, Mexico
JN539026.1, Hypersaline microbial mat, Guerrero Negro - Baja California Sur, Mexico
FN667354, Compost - Lahti, Finland
FN396772, Marine surface sediment, 50°C incubation - Smeerenburgfjorden, Svalbard
KM823684, River sediment - China
JQ407283, Mud volcano, Lei-gong-huo - eastern Taiwan
HQ916619, Mud volcano, Lei-gong-huo - eastern Taiwan
HQ916607, Mud volcano, Lei-gong-huo - eastern Taiwan
KF758687, Deep sea water
KF964589, Wetland soil - Ebinur lake
JN539026, Hypersaline mat, Guerrero Negro - Baja California Sur, Mexico
JN537682, Hypersaline mat, Guerrero Negro - Baja California Sur, Mexico
JQ407274, Mud volcano, Lei-gong-huo - eastern Taiwan
FN356285, Produced water, 80°C in situ oil reservoir temperature - Dan and Halfdan oil fields, North Sea
FN356288, Produced water, 80°C in situ oil reservoir temperature - Dan and Halfdan oil fields, North Sea
FN356295, Produced water, 80°C in situ oil reservoir temperature - Dan and Halfdan oil fields, North Sea
DQ647144, Produced water, 70°C in situ oil field temperature - Troll Formation, North Sea
FN356335, Produced water, 80°C in situ oil reservoir temperature - Dan and Halfdan oil fields, North Sea
FN356327, Produced water, 80°C in situ oil reservoir temperature - Dan and Halfdan oil fields, North Sea
FN356332, Produced water, 80°C in situ oil reservoir temperature - Dan and Halfdan oil fields, North Sea
FN356342, Produced water, 80°C in situ oil reservoir temperature - Dan and Halfdan oil fields, North Sea
FN356328, Produced water, 80°C in situ oil reservoir temperature - Dan and Halfdan oil fields, North Sea
FN356302, Produced water, 80°C in situ oil reservoir temperature - Dan and Halfdan oil fields, North Sea
FN356340, Produced water, 80°C in situ oil reservoir temperature - Dan and Halfdan oil fields, North Sea
FN356237, Produced water, 80°C in situ oil reservoir temperature - Dan and Halfdan oil fields, North Sea
FN356323, Produced water, 80°C in situ oil reservoir temperature - Dan and Halfdan oil fields, North Sea
DQ647125, Produced water, 70°C in situ oil field temperature - Troll Formation, North Sea
FN356239, Produced water, 80°C in situ oil reservoir temperature - Dan and Halfdan oil fields, North Sea
FN356341, Produced water, 80°C in situ oil reservoir temperature - Dan and Halfdan oil fields, North Sea
FN356304, Produced water, 80°C in situ oil reservoir temperature - Dan and Halfdan oil fields, North Sea
FN356314, Produced water, 80°C in situ oil reservoir temperature - Dan and Halfdan oil fields, North Sea
FN356303, Produced water, 80°C in situ oil reservoir temperature - Dan and Halfdan oil fields, North Sea
X77837
GU118213, Coral - Caribbean
KC668910, Coral - Red Sea
KC668846.1, Coral - Red Sea
KC668846, Coral - Red Sea
KC668837, Coral - Red Sea
KP305708.1, Reef coral - Luhuitou fringing reef, China
HQ606285.1, Marine sediments - South China Sea
KT973497.1, Intertidal outcrops - Isla de Mona, Puerto Rico
GQ267134.1, Hydrothermal sediments - Mothra Field, Juan de Fuca Ridge
HQ696463.1, Deep sea sediment - Indian Ocean
**ASV41, *Paramaledivibacter***
FR695371.1, Marine sediment, 25°C incubation - Aarhus Bay, Denmark
DQ831102.1
**ASV2045, *Paramaledivibacter***
**ASV60, *Paramaledivibacter***
KX062021, Hot spring, Polichnitos - Lesvos, Greece
AF458779, *Paramaledivibacter caminithermalis*, Deep sea hydrothermal chimney - Atlantic Ocean Ridge
FJ203551.1, Coral - Caribbean
**ASV165, *Paramaledivibacter***
KC668883.1, Coral - Red Sea
KC668869.1, Coral - Red Sea
KC668910.1, Coral - Red Sea
EU573106, Produced water, 131°C in situ oil reservoir temperature - Ekofisk oil field, North Sea
HQ696463, Deep sea sediment - Indian Ocean
AB806232, Ocean drilling core - Shimokita Peninsula, Japan
FJ203551, Coral - Caribbean
FJ202390, Coral - Caribbean
JQ515752, Coral - Caribbean
KC631808, Hypersaline microbial mat - Kiribati
EF123532, Coral - Caribbean
DQ446118, Coral - Caribbean
JX391232, Surface marine sediment - Hong Kong, China
KC668891, Coral - Red Sea
KT783480, Wukongibacter baidiensis, Deep sea hydrothermal field - Southwest Indian Ridge
JMSU01000863, Marine intertidal flat - Wadden Sea, Germany
AB806231, Ocean drilling core - Shimokita Peninsula, Japan

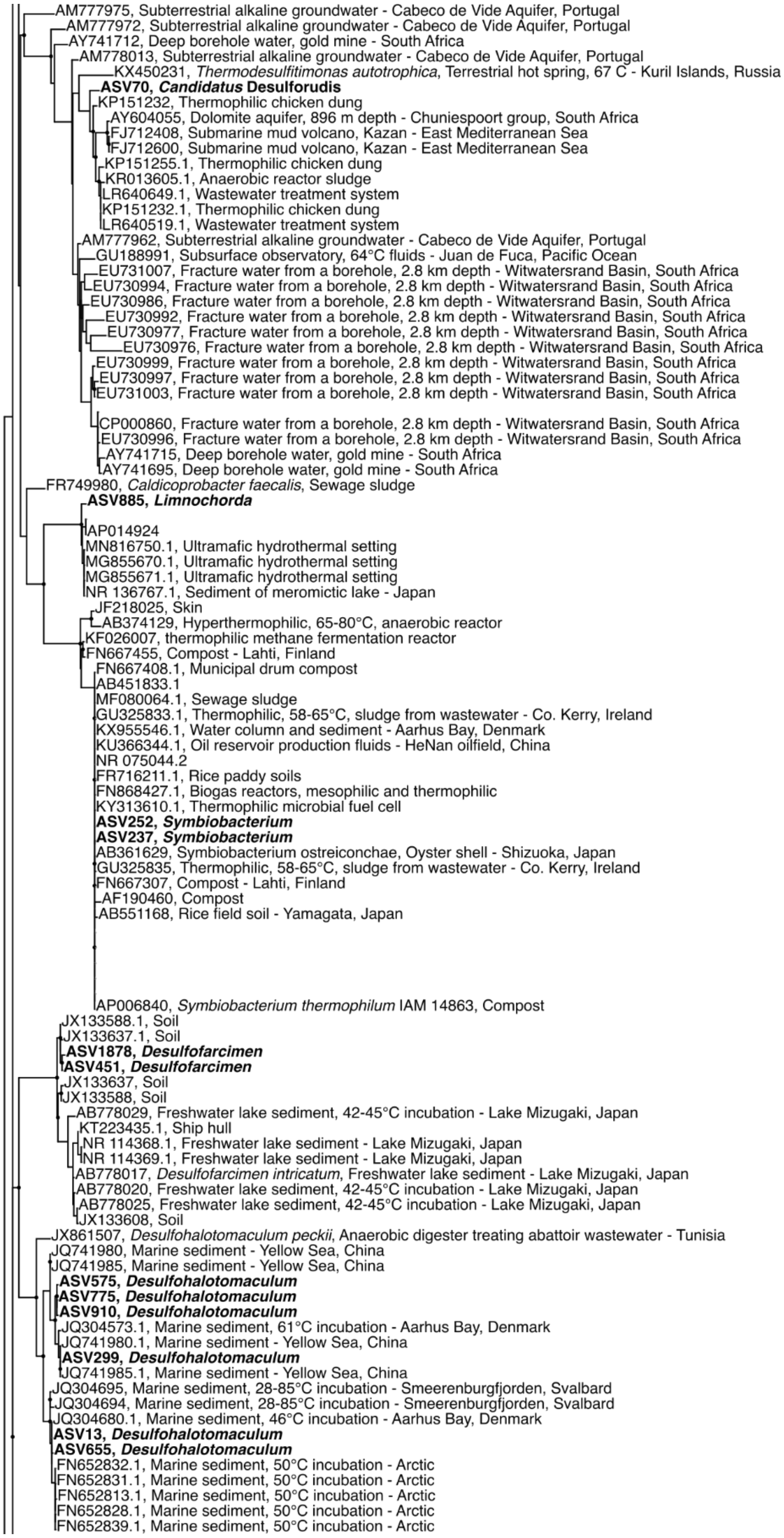
AM777975, Subterrestrial alkaline groundwater - Cabeco de Vide Aquifer, Portugal
AM777972, Subterrestrial alkaline groundwater - Cabeco de Vide Aquifer, Portugal
AY741712, Deep borehole water, gold mine - South Africa
AM778013, Subterrestrial alkaline groundwater - Cabeco de Vide Aquifer, Portugal
KX450231, *Thermodesulfitimonas autotrophica*, Terrestrial hot spring, 67 C - Kuril Islands, Russia
**ASV70, *Candidatus* Desulforudis**
KP151232, Thermophilic chicken dung
AY604055, Dolomite aquifer, 896 m depth - Chuniespoort group, South Africa
FJ712408, Submarine mud volcano, Kazan - East Mediterranean Sea
FJ712600, Submarine mud volcano, Kazan - East Mediterranean Sea
KP151255.1, Thermophilic chicken dung
KR013605.1, Anaerobic reactor sludge
LR640649.1, Wastewater treatment system
KP151232.1, Thermophilic chicken dung
LR640519.1, Wastewater treatment system
AM777962, Subterrestrial alkaline groundwater - Cabeco de Vide Aquifer, Portugal
GU188991, Subsurface observatory, 64°C fluids - Juan de Fuca, Pacific Ocean
EU731007, Fracture water from a borehole, 2.8 km depth - Witwatersrand Basin, South Africa
EU730994, Fracture water from a borehole, 2.8 km depth - Witwatersrand Basin, South Africa
EU730986, Fracture water from a borehole, 2.8 km depth - Witwatersrand Basin, South Africa
EU730992, Fracture water from a borehole, 2.8 km depth - Witwatersrand Basin, South Africa
EU730977, Fracture water from a borehole, 2.8 km depth - Witwatersrand Basin, South Africa
EU730976, Fracture water from a borehole, 2.8 km depth - Witwatersrand Basin, South Africa
EU730999, Fracture water from a borehole, 2.8 km depth - Witwatersrand Basin, South Africa
EU730997, Fracture water from a borehole, 2.8 km depth - Witwatersrand Basin, South Africa
EU731003, Fracture water from a borehole, 2.8 km depth - Witwatersrand Basin, South Africa
CP000860, Fracture water from a borehole, 2.8 km depth - Witwatersrand Basin, South Africa
EU730996, Fracture water from a borehole, 2.8 km depth - Witwatersrand Basin, South Africa
AY741715, Deep borehole water, gold mine - South Africa
AY741695, Deep borehole water, gold mine - South Africa
FR749980, *Caldicoprobacter faecalis*, Sewage sludge
**ASV885, *Limnochorda***
AP014924
MN816750.1, Ultramafic hydrothermal setting
MG855670.1, Ultramafic hydrothermal setting
MG855671.1, Ultramafic hydrothermal setting
NR 136767.1, Sediment of meromictic lake - Japan
JF218025, Skin
AB374129, Hyperthermophilic, 65-80°C, anaerobic reactor
KF026007, thermophilic methane fermentation reactor
FN667455, Compost - Lahti, Finland
FN667408.1, Municipal drum compost
AB451833.1
MF080064.1, Sewage sludge
GU325833.1, Thermophilic, 58-65°C, sludge from wastewater - Co. Kerry, Ireland
KX955546.1, Water column and sediment - Aarhus Bay, Denmark
KU366344.1, Oil reservoir production fluids - HeNan oilfield, China
NR 075044.2
FR716211.1, Rice paddy soils
FN868427.1, Biogas reactors, mesophilic and thermophilic
KY313610.1, Thermophilic microbial fuel cell
**ASV252, *Symbiobacterium***
**ASV237, *Symbiobacterium***
AB361629, Symbiobacterium ostreiconchae, Oyster shell - Shizuoka, Japan
GU325835, Thermophilic, 58-65°C, sludge from wastewater - Co. Kerry, Ireland
FN667307, Compost - Lahti, Finland
AF190460, Compost
AB551168, Rice field soil - Yamagata, Japan
AP006840, *Symbiobacterium thermophilum* IAM 14863, Compost
JX133588.1, Soil
JX133637.1, Soil
**ASV1878, *Desulfofarcimen***
**ASV451, *Desulfofarcimen***
JX133637, Soil
JX133588, Soil
AB778029, Freshwater lake sediment, 42-45°C incubation - Lake Mizugaki, Japan
KT223435.1, Ship hull
NR 114368.1, Freshwater lake sediment - Lake Mizugaki, Japan
NR 114369.1, Freshwater lake sediment - Lake Mizugaki, Japan
AB778017, *Desulfofarcimen intricatum*, Freshwater lake sediment - Lake Mizugaki, Japan
AB778020, Freshwater lake sediment, 42-45°C incubation - Lake Mizugaki, Japan
AB778025, Freshwater lake sediment, 42-45°C incubation - Lake Mizugaki, Japan
JX133608, Soil
JX861507, *Desulfohalotomaculum peckii*, Anaerobic digester treating abattoir wastewater - Tunisia
JQ741980, Marine sediment - Yellow Sea, China
JQ741985, Marine sediment - Yellow Sea, China
**ASV575, *Desulfohalotomaculum***
**ASV775, *Desulfohalotomaculum***
**ASV910, *Desulfohalotomaculum***
JQ304573.1, Marine sediment, 61°C incubation - Aarhus Bay, Denmark
JQ741980.1, Marine sediment - Yellow Sea, China
**ASV299, *Desulfohalotomaculum***
JQ741985.1, Marine sediment - Yellow Sea, China
JQ304695, Marine sediment, 28-85°C incubation - Smeerenburgfjorden, Svalbard
JQ304694, Marine sediment, 28-85°C incubation - Smeerenburgfjorden, Svalbard
JQ304680.1, Marine sediment, 46°C incubation - Aarhus Bay, Denmark
**ASV13, *Desulfohalotomaculum***
**ASV655, *Desulfohalotomaculum***
FN652832.1, Marine sediment, 50°C incubation - Arctic
FN652831.1, Marine sediment, 50°C incubation - Arctic
FN652813.1, Marine sediment, 50°C incubation - Arctic
FN652828.1, Marine sediment, 50°C incubation - Arctic
FN652839.1, Marine sediment, 50°C incubation - Arctic

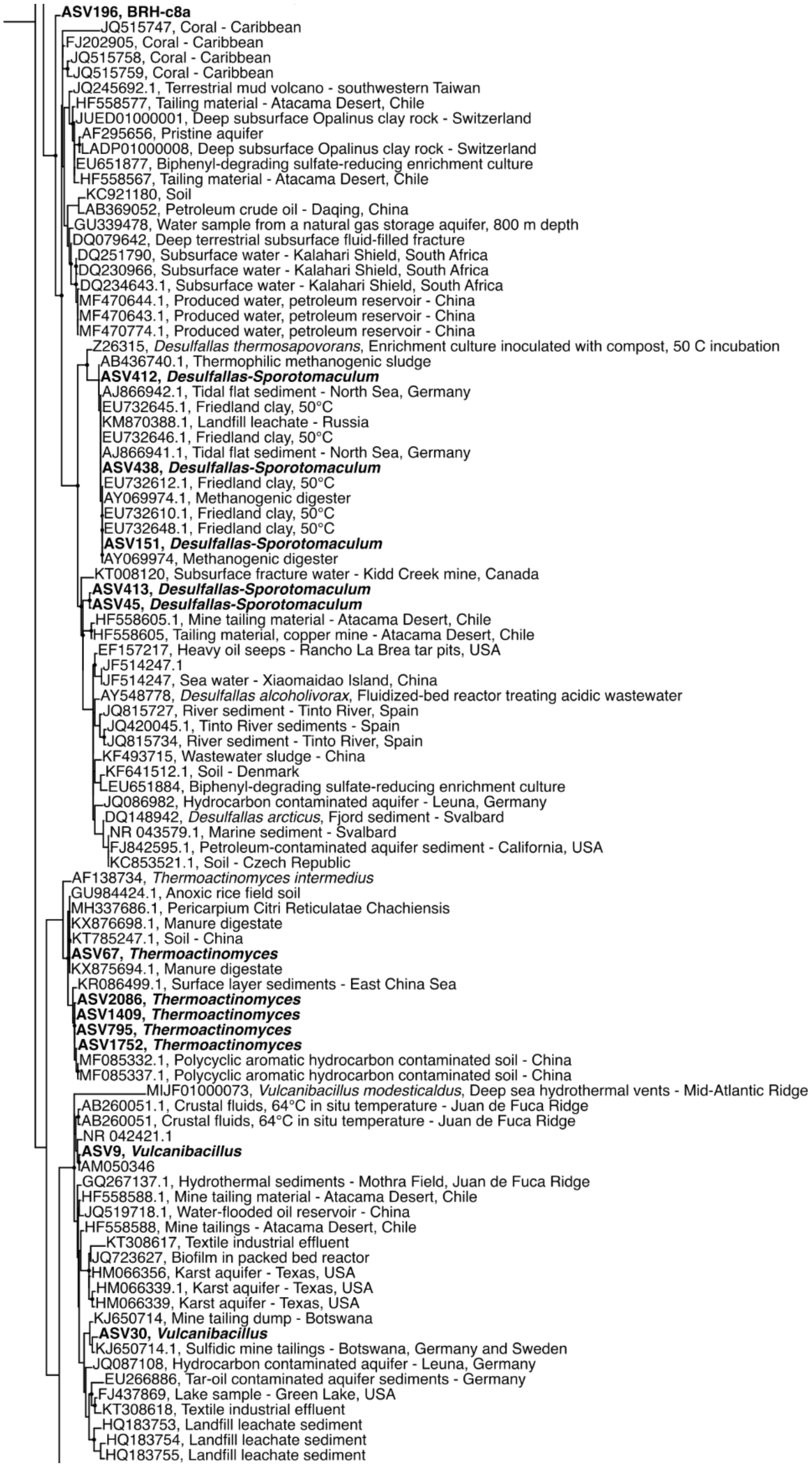
ASV196, BRH-c8a
JQ515747, Coral - Caribbean
FJ202905, Coral - Caribbean
JQ515758, Coral - Caribbean
JQ515759, Coral - Caribbean
JQ245692.1, Terrestrial mud volcano - southwestern Taiwan
HF558577, Tailing material - Atacama Desert, Chile
JUED01000001, Deep subsurface Opalinus clay rock - Switzerland
AF295656, Pristine aquifer
LADP01000008, Deep subsurface Opalinus clay rock - Switzerland
EU651877, Biphenyl-degrading sulfate-reducing enrichment culture
HF558567, Tailing material - Atacama Desert, Chile
KC921180, Soil
AB369052, Petroleum crude oil - Daqing, China
GU339478, Water sample from a natural gas storage aquifer, 800 m depth
DQ079642, Deep terrestrial subsurface fluid-filled fracture
DQ251790, Subsurface water - Kalahari Shield, South Africa
DQ230966, Subsurface water - Kalahari Shield, South Africa
DQ234643.1, Subsurface water - Kalahari Shield, South Africa
MF470644.1, Produced water, petroleum reservoir - China
MF470643.1, Produced water, petroleum reservoir - China
MF470774.1, Produced water, petroleum reservoir - China
Z26315, Desulfallas thermosapovorans, Enrichment culture inoculated with compost, 50 C incubation
AB436740.1, Thermophilic methanogenic sludge
ASV412, Desulfallas-Sporotomaculum
AJ866942.1, Tidal flat sediment - North Sea, Germany
EU732645.1, Friedland clay, 50°C
KM870388.1, Landfill leachate - Russia
EU732646.1, Friedland clay, 50°C
AJ866941.1, Tidal flat sediment - North Sea, Germany
ASV438, Desulfallas-Sporotomaculum
EU732612.1, Friedland clay, 50°C
AY069974.1, Methanogenic digester
EU732610.1, Friedland clay, 50°C
EU732648.1, Friedland clay, 50°C
ASV151, Desulfallas-Sporotomaculum
AY069974, Methanogenic digester
KT008120, Subsurface fracture water - Kidd Creek mine, Canada
ASV413, Desulfallas-Sporotomaculum
ASV45, Desulfallas-Sporotomaculum
HF558605.1, Mine tailing material - Atacama Desert, Chile
HF558605, Tailing material, copper mine - Atacama Desert, Chile
EF157217, Heavy oil seeps - Rancho La Brea tar pits, USA
JF514247.1
JF514247, Sea water - Xiaomaidao Island, China
AY548778, Desulfallas alcoholivorax, Fluidized-bed reactor treating acidic wastewater
JQ815727, River sediment - Tinto River, Spain
JQ420045.1, Tinto River sediments - Spain
JQ815734, River sediment - Tinto River, Spain
KF493715, Wastewater sludge - China
KF641512.1, Soil - Denmark
EU651884, Biphenyl-degrading sulfate-reducing enrichment culture
JQ086982, Hydrocarbon contaminated aquifer - Leuna, Germany
DQ148942, Desulfallas arcticus, Fjord sediment - Svalbard
NR 043579.1, Marine sediment - Svalbard
FJ842595.1, Petroleum-contaminated aquifer sediment - California, USA
KC853521.1, Soil - Czech Republic
AF138734, Thermoactinomyces intermedius
GU984424.1, Anoxic rice field soil
MH337686.1, Pericarpium Citri Reticulatae Chachiensis
KX876698.1, Manure digestate
KT785247.1, Soil - China
ASV67, Thermoactinomyces
KX875694.1, Manure digestate
KR086499.1, Surface layer sediments - East China Sea
ASV2086, Thermoactinomyces
ASV1409, Thermoactinomyces
ASV795, Thermoactinomyces
ASV1752, Thermoactinomyces
MF085332.1, Polycyclic aromatic hydrocarbon contaminated soil - China
MF085337.1, Polycyclic aromatic hydrocarbon contaminated soil - China
MIJF01000073, Vulcanibacillus modesticaldus, Deep sea hydrothermal vents - Mid-Atlantic Ridge
AB260051.1, Crustal fluids, 64°C in situ temperature - Juan de Fuca Ridge
AB260051, Crustal fluids, 64°C in situ temperature - Juan de Fuca Ridge
NR 042421.1
ASV9, Vulcanibacillus
AM050346
GQ267137.1, Hydrothermal sediments - Mothra Field, Juan de Fuca Ridge
HF558588.1, Mine tailing material - Atacama Desert, Chile
JQ519718.1, Water-flooded oil reservoir - China
HF558588, Mine tailings - Atacama Desert, Chile
KT308617, Textile industrial effluent
JQ723627, Biofilm in packed bed reactor
HM066356, Karst aquifer - Texas, USA
HM066339.1, Karst aquifer - Texas, USA
HM066339, Karst aquifer - Texas, USA
KJ650714, Mine tailing dump - Botswana
ASV30, Vulcanibacillus
KJ650714.1, Sulfidic mine tailings - Botswana, Germany and Sweden
JQ087108, Hydrocarbon contaminated aquifer - Leuna, Germany
EU266886, Tar-oil contaminated aquifer sediments - Germany
FJ437869, Lake sample - Green Lake, USA
KT308618, Textile industrial effluent
HQ183753, Landfill leachate sediment
HQ183754, Landfill leachate sediment
HQ183755, Landfill leachate sediment

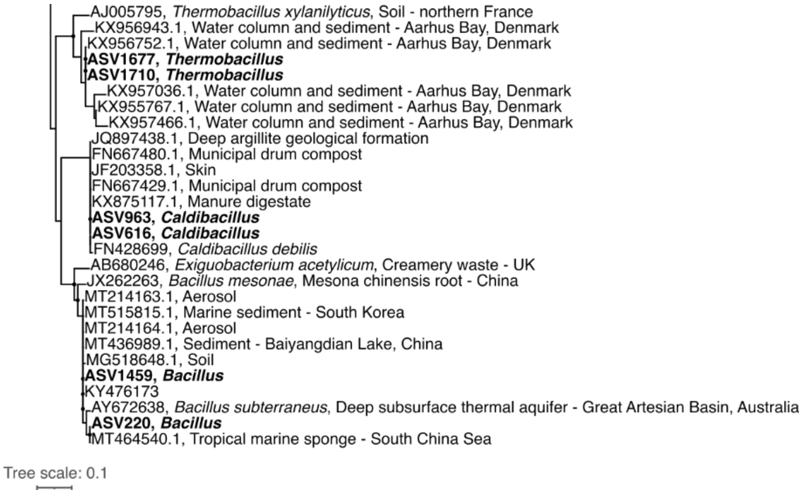


**Extended Data Fig. 4 | Phylogenetic association of seep-associated sequences with sequences from other environments.** Maximum likelihood tree showing phylogenetic relationships between 42 seep-associated ASVs (bold) and close relatives in the GenBank database. Black circles at the branch nodes indicate >80% bootstrap support (1,000 re-samplings). Scale bar indicates 10% sequence divergence as inferred from PhyML. *Pseudomonas aeruginosa* (accession number Z76672) was used as an outgroup to root the tree.

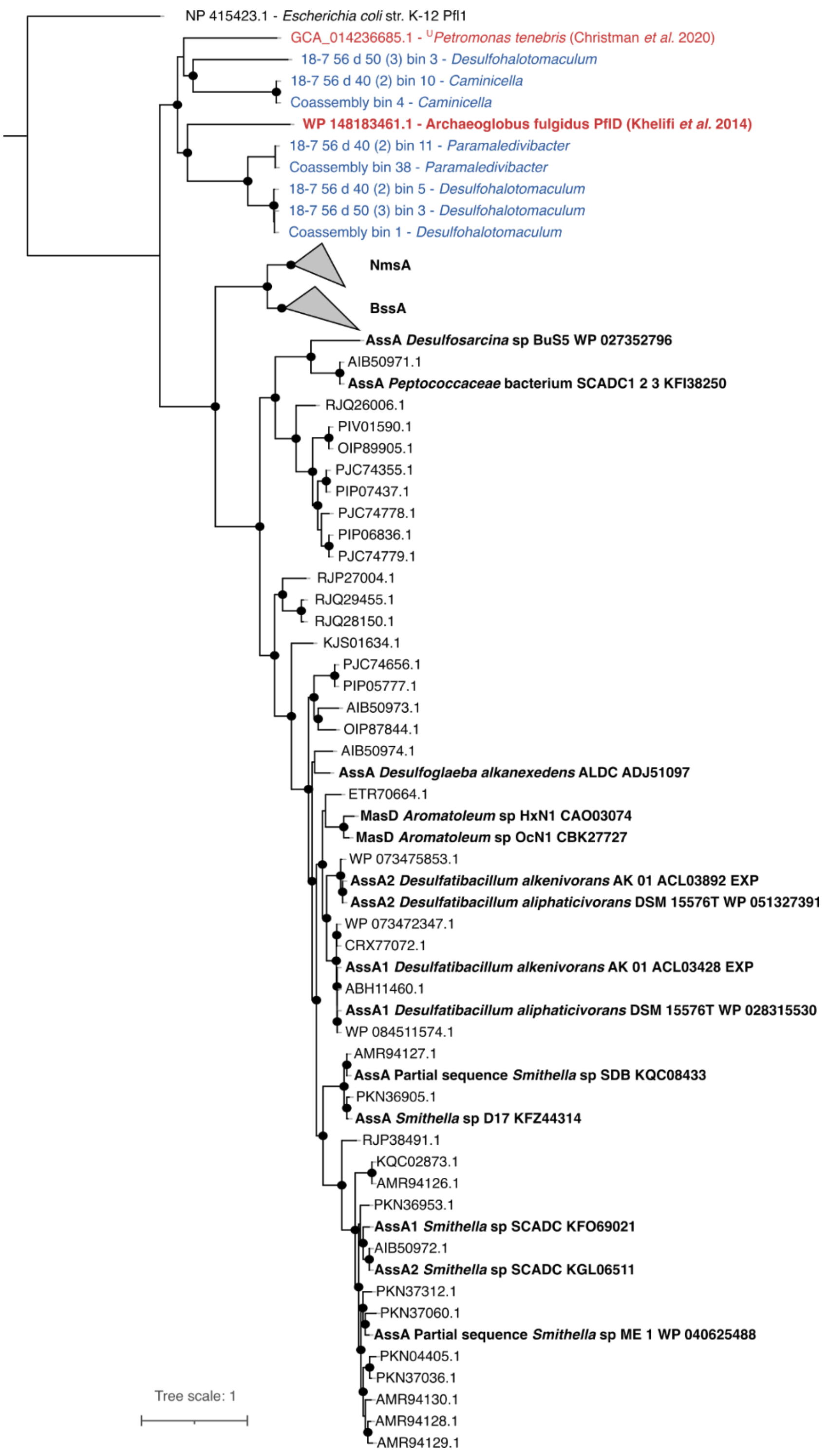


**Extended Data Fig. 5 | Phylogenetic relationships of putative glycyl-radical enzymes with alkylsuccinate synthases.** Putative anaerobic alkane-degrading pyruvate-formate lyase enzyme variants from *Desulfohalotomaculum*, *Caminicella* and *Parameldivibacter* thermophilic spores in this study (shown in blue) cluster together with homologous *pflD* gene sequences found in the oil reservoir bacteria $^{U}$*Petromonas tenebris*[34] and the oil reservoir archaea *Archaeoglobus fulgidus* VC-16, shown in red, that can degrade alkanes at high temperature under anaerobic conditions[35]. Reference sequences of alkane succinate synthase (AssA and MasD) genes with corresponding experimental verification of anaerobic alkane degradation are shown in bold. Benzyl succinate synthase (BssA) and naphthyl-2-methyl-succinate synthase (NmsA) sequences are represented by collapsed clades. Black circles at the branch nodes indicate >80% bootstrap support (1,000 re-samplings). Scale bar indicates 10% sequence divergence as inferred from PhyML. A sequence of pyruvate formate lyase (Pfl) from *E. coli* was used to root the tree.

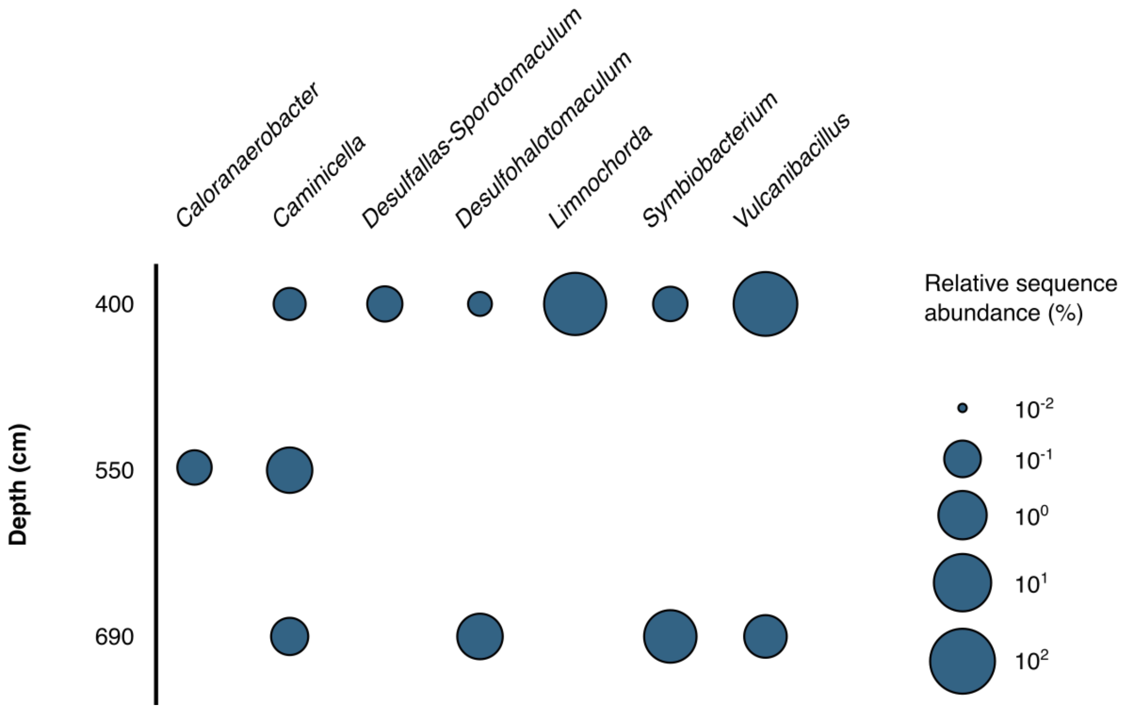


**Extended Data Fig. 6 | Endospores remain viable during burial.** Detection of endospore-forming bacteria with the same taxonomic classification as seep-associated ASVs (c.f. Supplementary Table 5) in deeper Scotian Slope sediment layers following incubation at 50°C. Bubble size indicates the maximum relative sequence abundance of a taxon in unrarefied sequence libraries corresponding to the indicated depth.
